## Supporting info (SI) for "Temperature-triggered *in situ* forming lipid mesophase gel for local treatment of ulcerative colitis"

**Small angle X-ray scattering**

Small‐angle X-ray scattering (SAXS) was used to determine the lipid phase, and, thus, construct the partial phase diagram, for different MLO–water systems. That method relies on constructive interferences in the reciprocal space from many ordered scattering planes that belong to the mesophase. An X-ray beam is directed at the lipid sample, and the resulting scattering pattern gives a characteristic set of rings, or maxima, that correspond to Bragg reflections. Their positions in the reciprocal space depend on the Miller indices of the mesophase scattering planes, and the sequence of Bragg reflections (and their ratio) consequently identifies the symmetry of the mesophase studied. SAXS allows for the lattice parameter—the size of the repeat unit cell—to be determined. When the parameter is translated, researchers can reconstruct the entire mesophase in 3D. SAXS measurements were used to determine the phase identity and symmetry of the produced LMPs. Measurements were performed on a Bruker AXS Micro, as described in the main text. MLO was used as the lipid component of the mesophases and mixed with weighed amounts of drugs (10% w/w) in sealed Pyrex tubes and alternatively centrifuging (10 min, 5000 g) several times at room temperature until a homogenous mixture was obtained. The mesophase was then equilibrated for 48 h at room temperature in the dark.

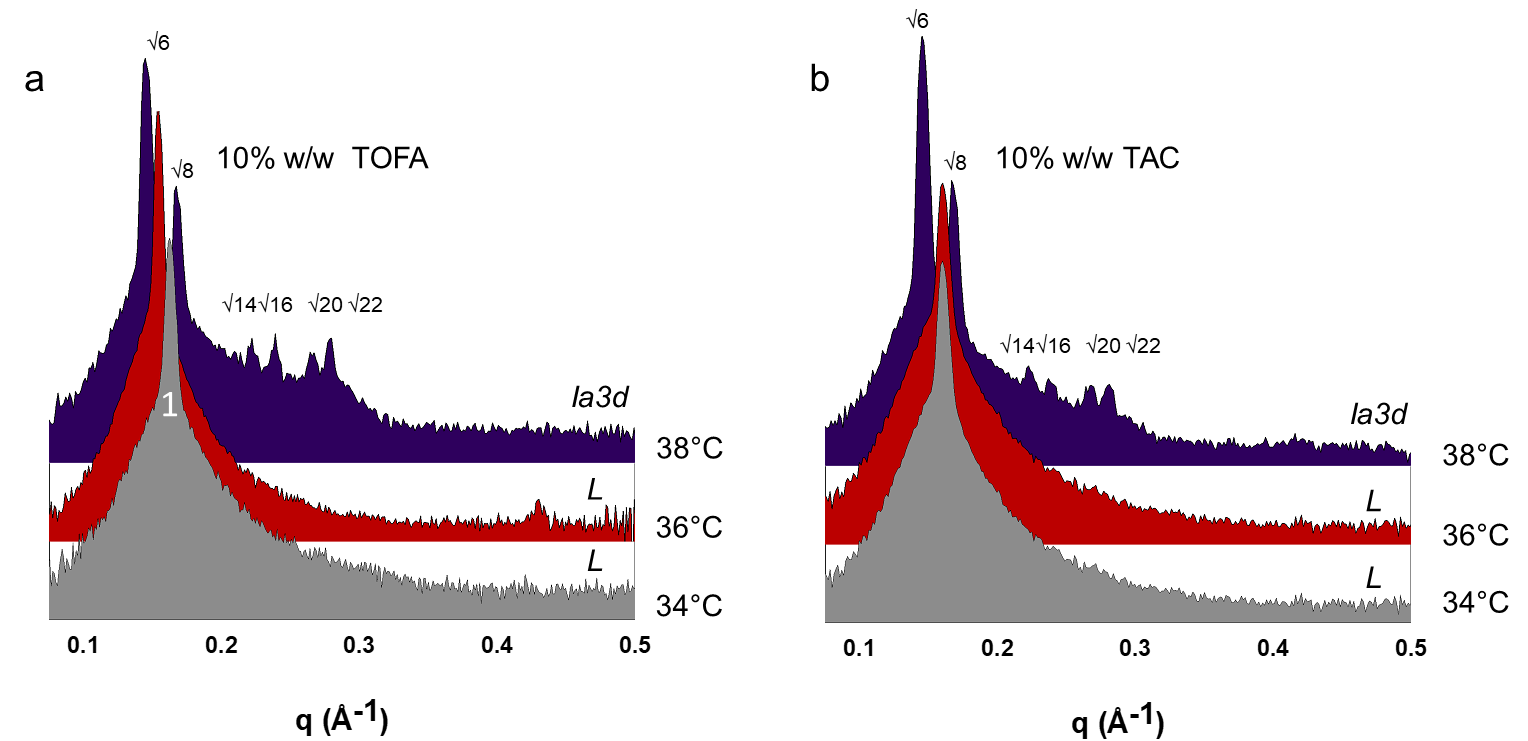

**Figure S1.** SAXS spectra acquired at different temperatures on gels containing 10% w/w of TOFA (a) and 10% w/w of TAC (b).

**HPLC method: Tofacitinib Citrate**

Tofacitinib citrate (TOFA) was detected by reverse-phase liquid chromatography using a Macherey-Nagel Nucleosil 100-5 C18 (4.0 x 250 mm; 5.0 µm particle size) column. The mobile phase consisted of acetonitrile/methanol/water (13:13:74 v/v) + 0.1% trifluoroacetic acid at a flow rate of 1 mL/min, temperature 25 °C and UV detection at λ = 278 nm. An internal standard (caffeine, 20 µg/mL) was added to each sample to correct for inter-injection variation and UV detection at λ = 278 nm. Data were collected and analyzed using the software Chromeleon 7 (Thermo Fisher).

**HPLC method: Tacrolimus**

Tacrolimus (TAC) was detected by reverse-phase liquid chromatography using a Macherey-Nagel Nucleosil 100-5 C18 (4.0 x 250 mm; 5.0 µm particle size) column. The mobile phase consisted of methanol/water (80:20 v/v) + 0.1% trifluoroacetic acid at a flow rate of 1 mL/min, temperature 50 °C and UV detection at λ = 214 nm. An internal standard (ketoconazole, 20 µg/mL) was added to each sample to correct for inter-injection variation and UV detection at λ = 278 nm. Data were collected and analyzed using the software Chromeleon 7 (Thermo Fisher).

**Stability Study of TAC and TOFA**

The stability of the drugs – TOFA and TAC – was monitored over one month. At specific time points, an aliquot of the formulation was analyzed at the HPLC, and the content of the drug recorded. Data are expressed as relative percentage referred to day 0.

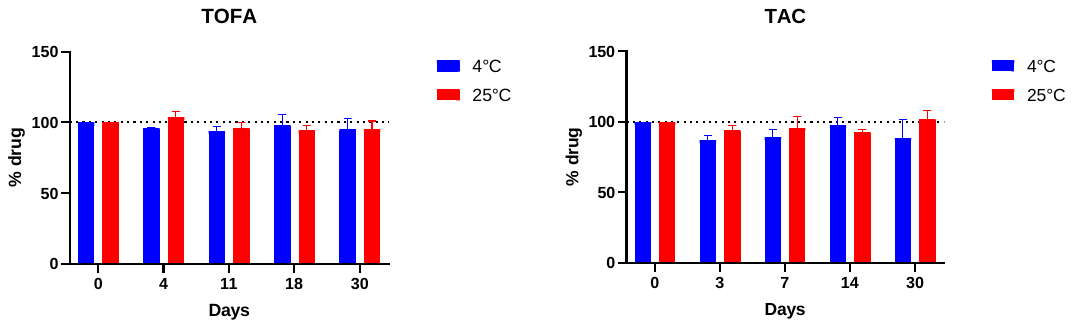

**Figure S2.** Long-term stability of TOFA loaded into TIF-gel and TAC loaded into TIF-gel over one month. Data are expressed as percentage ± SD.

**Dead Volume of the syringe and canula**

To calculate the dead volume of the syringe 1 ml: Injekt®-F (Fine Dosage) Luer Solo (Luer Slip) (Braun) with the canula (size 20G, L × diam. 1.5 in. × 1.9 mm ), the syringe was filled with different amount of formulation Monolinolein + MilliQ water (84% lipid and 16% water) and the amount that came out of the syringe was recorded.

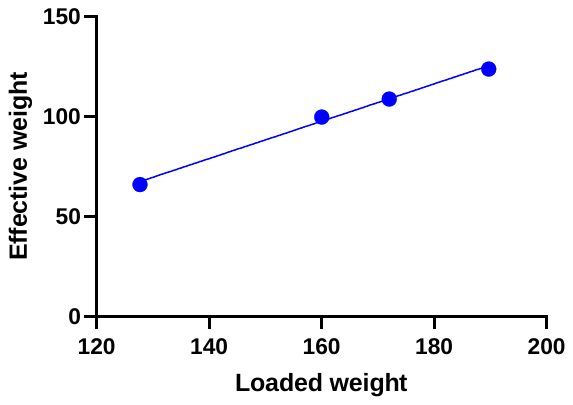

**Figure S3**. Calibration curve used to calculate the amount of formulation that has to be loaded into the syringe to obtain exactly 100 mg of formulation from the syringe with the canula.

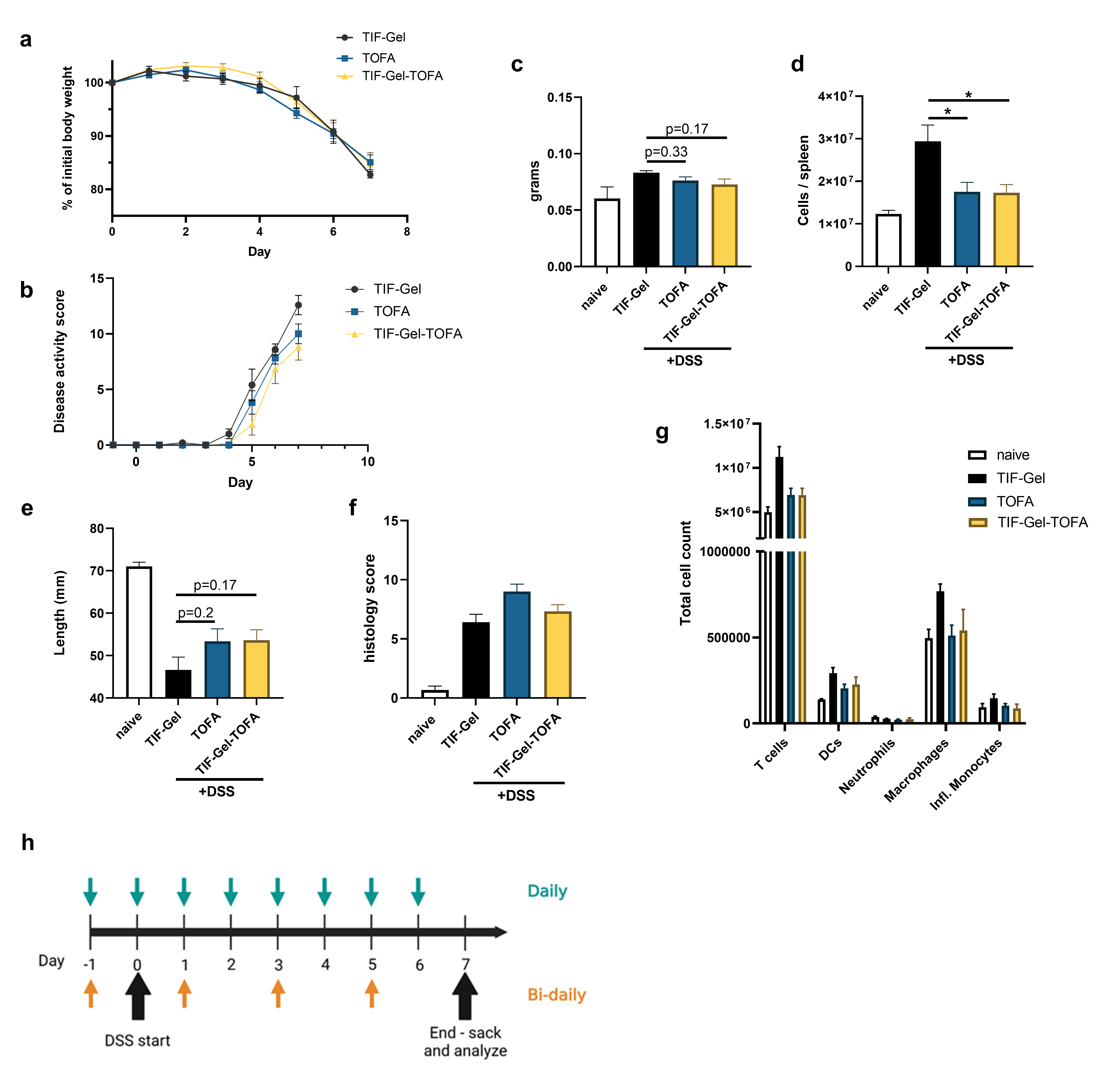

**Figure S4: Daily application of TIF-Gel-TOFA did not improve animal health compared to TOFA in vehicle.** Mice were treated rectally with empty gel (TIF-gel), TOFA in vehicle (TOFA) or drug-loaded gel (TIF-Gel-TOFA) daily from 1 day before the start of DSS treatment until day 6). During the treatment, mice were weighed (a), and the severity of their illness was assessed (b). At day 7, mice were euthanized, and various disease parameters were recorded included spleen weight and cellularity (c and d), colon length and pathology (e and f), and the total populations of various immune cells from spleens were calculated (g). h) An application scheme detailing the timeline for the daily and bi-daily rectal applications of the various compounds. Statistical values were calculated by one-way ANOVA (c, d, e, f), multiple T tests per group with Holm-Sidak correction (g) or two-way ANOVA (a and b). *: p<0.05, where no value is indicated, p>0.05). Naïve values were excluded from analyses. Error bars are ± SEM.

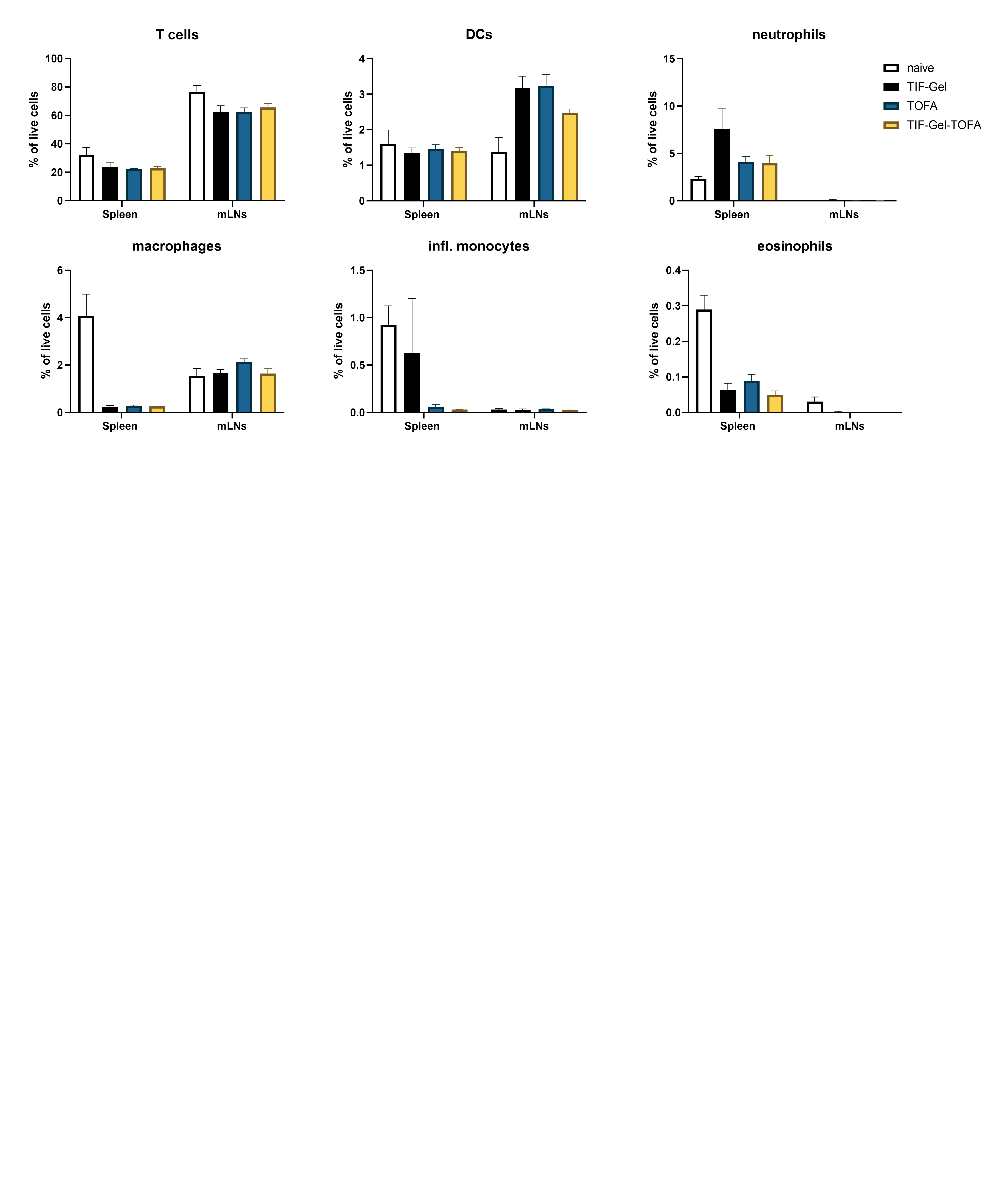

**Figure S5: DSS challenge TIF-gel-TOFA does not rescue the changes to cell frequencies of spleen and mLN in DSS-treated mice**. The relative numbers of various cells types in mouse spleens and mesenteric lymph nodes (mLNs) were quantified. No statistically significant differences in percentages were observed between the different treatment groups. Data were analyzed using one-way ANOVAs. DCs, dendritic cells. Error bars are ± SEM.
